# Profiles of Aging Based on Cognition, Affect, and Brain Reserve

**DOI:** 10.64898/2025.12.04.692428

**Authors:** Setthanan Jarukasemkit, Lyn Stahl, Karen M Tam, Bibo Feng, Samuel Naranjo Rincon, Xiaoke Luo, Hailey Modi, Kassandra Hamilton, Petra Lenzini, Fyzeen Ahmad, Ty Easley, Janine D Bijsterbosch

## Abstract

The aging paradox describes improvements in emotional wellbeing as a function of aging, despite declines in cognition. Conversely, late life depression has been associated with increased cognitive decline in aging. We sought to understand these seemingly contradictory patterns of cognitive and mental health in older age. Building on cognitive reserve, affective reserve, and brain reserve models of aging, we developed three alternative algorithmic approaches to group N=22,686 participants from the UK Biobank into different profiles of aging. Our results revealed that aging profiles identified using our data-driven brain reserve model, which incorporated measures of cognition, neuroticism, and brain volume, achieved the highest validation results. Importantly, only two of the four aging profiles were characterized by the aging paradox (i.e., improved emotionality and decreased cognition with age). We identified one profile characterized by particularly low levels of neuroticism and relative resilience to cognitive decline. Another profile benefited from relatively preserved brain volumes, potentially driven by younger ages and/or higher socioeconomic status. Conversely, we identified two profiles with poorer health characteristics, including one profile with elevated cardiovascular risk. Taken together, these findings enrich our understanding of the emotion paradox and highlight the value of taking a nuanced and stratified approach when studying aging. In the future, aging profiles could be used to target preventative strategies to address modifiable risk factors and improve lifespan and healthspan.

## 1. Introduction

The relationship between cognitive and mental health in older age is poorly understood. Cognition declines with age across domains (e.g., processing speed, memory, and reasoning) [1]. Conversely, emotional wellbeing improves with age [2], leading to reduced incidence of major depressive disorder consequent to normal aging [3]. The so-called ‘emotion paradox’ describes this apparent disconnect between improved emotional wellbeing and declines in cognition and physical health typical of aging [4]. Although depression incidence decreases with age, affective disorders remain a concern. Late-life depression is a risk factor for cognitive decline and dementia [5–7], suggesting a positive (rather than inverse) association between mental and cognitive health in aging. The goal of this paper was to disentangle this dual relationship between cognitive and mental health using a large-scale older aged cohort.

A possible explanation for these contradictory relationships may be different aging profiles [8,9]. As they age, individuals undergo varying cognitive and affective trajectories, with some proving less vulnerable to decline than others. Cognitive reserve describes resilience where cognitive abilities are less vulnerable to degradation [10–13]. Affective reserve describes emotional resilience; emotional symptoms remain relatively low despite adverse life circumstances or pathologies [14–16]. Cognitive and affective reserve may not be independent. Prior work revealed decreased apathy - but not depressed mood - as a function of increased cognitive reserve in older individuals [17], such that increased cognitive reserve may help mitigate negative emotional states. We defined aging profiles based on cognitive and affective reserve to assess potentially differential relationships between cognitive and mental health in relation to age.

In addition to cognitive and affective reserve, the brain reserve model offers a biologically-grounded theory to explain differences in aging. Brain reserve posits that differences in underlying structural neurobiology explain differential resistance to brain aging or pathology [18]. Higher brain reserve has been linked to lower mortality [19], reduced risk of Alzheimer’s Disease [20], and reduced risk of depressive episodes in late life [11]. As such, brain reserve may form a link between cognitive and affective reserve that may determine differential profiles of aging and help explain the emotion paradox. In this study, we assessed whether inclusion of brain reserve improved the determination of different profiles of aging.

To our knowledge, there is minimal research studying aging profiles based jointly on cognitive, affective, and brain reserve. To address this, we defined aging subtypes in an older-age population cohort to characterize different profiles of aging. We hypothesized the presence of four profiles of aging, namely: healthy aging (high cognitive and affective reserve), suboptimal aging (low cognitive and affective reserve), cognitive reserve (high cognitive but low affective reserve), and affective reserve (high affective but low cognitive reserve). We tested different analytical models to define aging profiles, assessed how many profiles of aging may exist, and determined demographic, neuroanatomical, and etiological differences between aging profiles. To determine if aging profiles contribute to the emotion paradox, we investigated profile-specific associations between age and cognition/ affect. Our results support the existence of aging profiles that differ in terms of the association between age and cognition/affect. These results inform our understanding of the emotion paradox and highlight the value of taking a nuanced and stratified approach to aging.

## 2. Materials and Methods

### 2.1 Dataset

Analyses were performed using data from the UK Biobank under application number 47267. Further details on the dataset [21], imaging acquisition [22], and preprocessing [23] can be found elsewhere. The overall combined sample included 22,686 subjects (mean age 64 +/-7.5 years, percent male 48.1%; Table 1) with complete data for cognition, affect, and brain reserve variables detailed below.

**Table 1.** Demographic Data

| Variable | UKB Variable IDs | Variable | Value |
| --- | --- | --- | --- |
| Sample Size | NA | NA | 22,686 |
| Age | 21003 | Integer | Mean = 64, SD = 7.545 |
| Percentage of Male | 31 | Categorical | 48.1% |
| Income | 738 | Categorical | Median = 3 (£31,000-£51,999), IQR = 2 |
| Neuroticism | Sum [27] of: 1920, 1930, 1940, 1950, 1960, 1970, 1980, 1990, 2000, 2010, 2020, 2030 | Categorical | Median = 4, IQR = 4 |
| Cognition | Weighted sum [24] of: 2016, 2023 | Continuous | Mean = 0.002, SD = 1.169 |

### 2.2 Variables to define aging profiles

Cognition was measured using the Lyall Model 3 composite score [24], which incorporates measures of verbal-numerical reasoning and log reaction time (Table 1). Negative emotionality was measured using the Eysenck Neuroticism score (N-12; Table 1), a twelve question measure assessing neuroticism domains [25–27]. Although this scale captures risk of psychopathology well [28,29], we note that it focuses on negative emotionality and may overlook positive emotional states. This study thus focuses on negative emotionality when assessing affect. The cognitive and negative emotionality measures were z-scored before analysis. Brain reserve was defined using multivariate measures of 81 gray matter volumes generated by freesurfer for DKT cortical and ASEG subcortical parcellations (Supplementary Table 1).

### 2.3 Defining aging profiles

We used three models to define aging profiles:

#### Predefined behavior model

Subjects were divided into quadrants based on cognitive and neuroticism scores. For both cognition and neuroticism, participants were assigned to high or low if they were above or below the 50th percentile, respectively (Table 1). Healthy aging was defined as high cognition/low neuroticism, affective reserve as low cognition/low neuroticism, cognitive reserve as high cognition/high neuroticism, and suboptimal aging as low cognition/high neuroticism. The advantage of this model is interpretability and straightforward application.

#### Data-driven behavior model

This model used identical summary variables for cognition and negative emotionality as the predefined model. Scores were entered into a fuzzy C-means clustering algorithm to define aging profiles. In addition to the hypothesized 4-profile solution, we tested the consistency of findings when changing the number of clusters/profiles to 2, 3, and 5. Fuzzy C-means initially assigns cluster membership randomly, and adjusts each point’s estimated cluster until the algorithm stabilizes at the error rate specified. Unlike hard clustering approaches, fuzzy C-means reports the degree to which a subject belongs to each cluster, rather than reporting one cluster assignment. Fuzzy C-means consequently offers an alternative to K-means clustering that determines the probability of cluster assignments for each participant. These probabilities can be used to assess uncertainty of cluster assignments through cluster entropy. Fuzzy c-means were calculated using the python package skfuzzy, with neuroticism and Lyall cognitive score as inputs. Error was set to 0.005 and maximum iterations to 1000.

#### Data-driven brain reserve model

The brain reserve model incorporated 81 gray matter volumes from 62 cortical and 19 subcortical regions (Supplementary Table 1). First, canonical correlation analysis (CCA) was performed to link gray matter volumes to summary variables (the cognitive and emotionality measures). Briefly, CCA finds the linear combination of sets of variables that maximize correlation between canonical brain variables (U) and canonical behavioral variables (V). The significance of CCA was tested by permuting the rows of the behavioral inputs 1,000 times while holding brain inputs fixed. For each permutation we recomputed CCA, and p-values were estimated by comparing the first canonical correlation as max-statistic against all permuted canonical correlations, providing family-wise error control over two multiple comparisons for the two canonical covariates. Both U and V variables from all significant canonical variables were used as inputs to the fuzzy C-means clustering algorithm as described above for each cluster/profile solution to test stability. We leveraged traditional CCA without regularization because our analysis included a sufficient subjects-to-feature ratio (273 to 1) to achieve high power and accuracy without regularization [30].

#### Effects of age

Although cognition and negative emotionality vary with age, we purposefully did not control for age prior to the definition of aging profiles. A core part of our analysis was investigating associations between age and cognition/negative emotionality separately within each aging profile. Such analyses would not be possible if we regressed age out prior to the definition of aging profiles. Furthermore, regressing out age at the outset risks collider effects and other statistical interactions that complicate interpretability. Although this approach comes with the caveat that profiles may differ on age, we felt the benefits outweighed this tradeoff. Age and sex were included as covariates in subsequent analyses to compare aging profiles.

### 2.4 Reliability of aging profiles by number of clusters

To assess model reliability, bootstrapping was performed by subsampling 80% of participants without replacement across 1000 bootstraps. These analyses were performed for behavior and brain-reserve models only because they require non-discrete cluster assignments. For comparison, we generated a null distribution by replacing fuzzy C-means inputs (matched in dimensionality) with random values from a normal distribution (mean=0, SD=1) and applied the same bootstrapping procedure. These comparisons against null data are important because clustering models will always derive a solution, which may achieve relatively high null performance depending on the clustering algorithm and dimensionality of the input values [31].

For each bootstrap, we estimated model stability using the Adjusted Rand Index (ARI) between the bootstrap cluster assignment and full-data cluster assignment as a measure of cluster stability. An ARI of 1 indicates identical/static assignments, whereas values near 0 indicate unstable cluster assignment. Additionally, we assessed cluster separability and cluster assignment overlap using silhouette scores and entropy, respectively. Silhouette scores were computed, where higher values indicate better separation and compactness of cluster. We were able to compute entropy as a measure of unstable cluster assignment due to the advantage of fuzzy C-means, where each subject (*i*)had cluster probabilities (*p*_*ic*_)across the total number of clusters (*p*_*i1*_*p*_*i2*,_ …,*p*_*ik*_). To compare across different numbers of clusters (_*k=2-5*_), we normalized entropy by *k*:

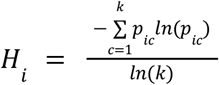

Differences between the data and null distributions were tested using a paired non-parametric bootstrap test. For each of 1,000 bootstrap resamples, we computed the reliability metric for the data and the corresponding null data, then calculated the difference between true results and null. We obtained an one-sided empirical p-value by counting how many bootstrap differences were < 0 for ARI and Silhouette scores, and > 0 for entropy, and dividing by the total number of resamples (1,000). With 1,000 bootstraps, the smallest attainable p-value was 0.001.

### 2.5 Contribution to the emotion paradox

To assess whether profiles of aging reflect predicted relationships between cognition, neuroticism, and age, we assessed correlations between these variables by calculating Spearman correlations separately in each identified aging profile, focusing on the four-profile solutions. We hypothesized that some profiles would capture the negative correlation between age-cognition and age-neuroticism predicted by the emotion paradox, and other profiles may reveal different associations between age and cognition/neuroticism capturing complex cognition-affect associations as a function of age.

A post hoc windowed analysis was subsequently performed to calculate Spearman correlations between age and cognition/neuroticism for 15-year windows in steps of 3 years. These window settings were chosen to ensure sufficient sample size (minimum N=1,000) in each window to accurately estimate associations. This analysis was performed to assess the stability of cross-sectional associations between age and cognition/neuroticism in each profile as a function of age.

### 2.6 Neuroanatomical differences between aging profiles

To visualize differences in regional brain volumes across aging profiles, we plotted the z-scored mean for each region within each aging profile. This allowed neuroanatomical characteristics of different aging profiles to be seen, highlighting which regions tend to be relatively larger or smaller in between groups. Notably, no statistical comparisons were performed to avoid circularity because gray matter volumes were used as inputs to the brain reserve model.

### 2.7 Validity of aging profiles by etiological relevancy

To assess whether aging profiles differed significantly, we statistically compared the resulting profiles on relevant dimensions of demographics, brain size, cardiovascular health, inflammation, and polygenic risk score for Alzheimer’s Disease. In total, 11 variables were tested as part of the validation analyses (Supplementary Table 2).

Statistical analyses were performed using analyses of variance (ANOVA). The main effect of aging profile (2-5 levels) was modeled separately for each validation variable, model solution, and as an analysis of covariance (ANCOVA) to allow covariate control for age and sex (and intracranial volume for white matter hyperintensities). Covariate-controlled analyses were not performed for age, sex, and total intracranial volume measures as these measures were used for covariates. Results were interpreted based on effect size (Ω), using standardized ranges of 0.01-0.06 for small effects, 0.06-0.14 for moderate effects, and 0.14> for large effects. Effect sizes were used instead of p-values to reduce the impact of sample size on each of the validation measures (range N=742 - 22,686) and to avoid overinterpretation of minimal (but significant) effects. Post-hoc Tukey-HSD tests were performed to determine which group differences contributed to the main effect. Tukey’s procedure controls family-wise error rate for pairwise comparisons within each validation variable.

## 3. Results

### 3.1 Aging profiles as defined using three different models

We extracted the four hypothesized profiles of aging using three different models (Fig. 1), and for dimensionalities from 2-5 profiles (Supplementary Fig. 1). The data-driven behavior solution was comparable to the predefined solution (ARI=0.56, Supplementary Fig. 2), identifying groups delineated by high/low cognitive scores and high/low negative emotionality. Notably, at high negative emotionality the data-driven behavior model preferentially placed individuals in the suboptimal aging group (profile 3).

**Figure 1.**
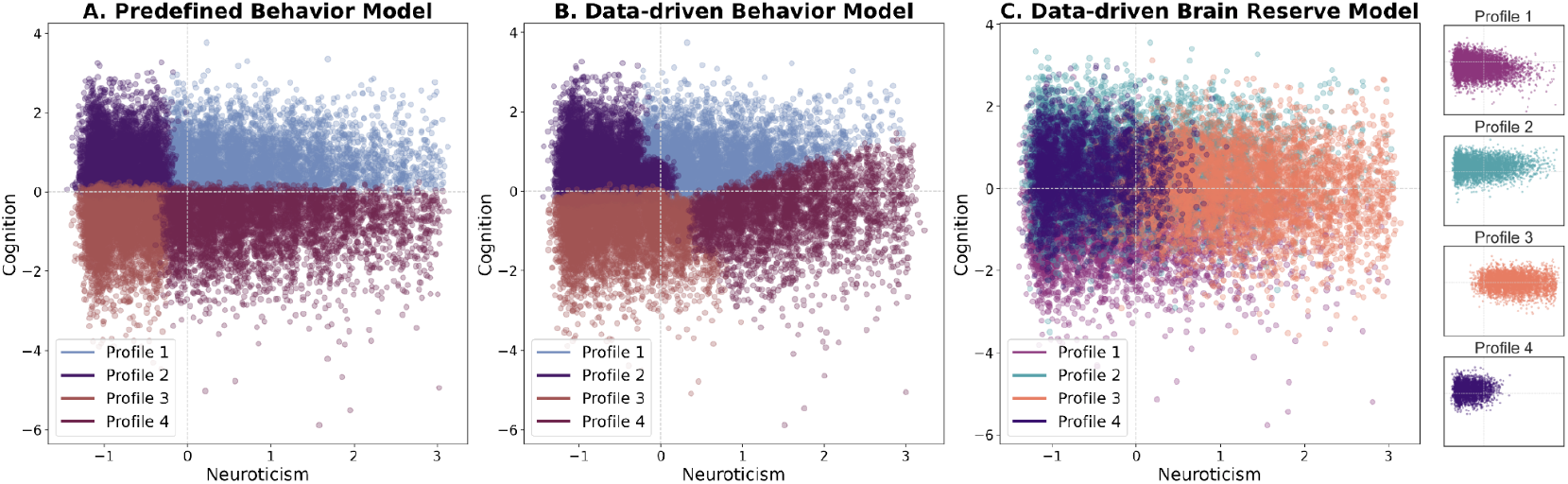
Four cluster solutions for each model. A) Predefined model profiles. B) Data-driven behavior model profiles. C) Data-driven brain reserve model profiles. A different color scheme was used for C because there is no clear one-to-one match of the brain reserve profiles to the other solutions. A minor amount of jitter was added to the data points to improve visualization of a large number of data points. Notably, the brain reserve profiles are shown in demeaned cognition-neuroticism space for direct visual comparison, yet clustering was performed in canonical covariate space explaining the observed overlap. Therefore, each brain reserve profile is also shown in a separate figure.

The brain reserve model differed from the predefined and data-driven behavior models (ARI=0.16 & 0.19, respectively; Supplementary Fig. 2). Cognitive score was the dominant variable used to determine group identity. This is likely driven by the close relationship between neural structural variables incorporated into the brain reserve model and cognitive score driving the first canonical component (r_U-V_=0.3047, p<0.001), compared to the second canonical component that was more associated with negative emotionality (r_U-V_=0.1590, p<0.001). The 4-profile solution in the brain reserve model did not clearly differentiate profiles in the space of cognitive and negative emotionality scores, which is expected given that clustering was performed in CCA space after extracting canonical components informed by composite scores as well as brain reserve. In CCA space, the canonical behavioral variables (V) varied as a function of cognition (V1) and neuroticism (V2), whereas the associated canonical brain variables (U) varied as a function of age (U1) and sex (U2; Supplementary Fig. 3). Notably, the brain reserve model placed a third cluster between and overlapping the healthy aging and affective reserve groups. Full demographic information regarding the 4 brain reserve profiles can be found in Supplementary Table 3.

We repeated the models (either based on only cognitive and affective reserve data or incorporating brain reserve data) across a number of different solutions (2, 3, 4, or 5 profiles, Supplementary Fig. 1). Interestingly, the 2-profile solution was highly unstable between the data-driven behavior and brain reserve models. This may be explained by correlation between the neural components of the model and cognition, leading to cognition explaining a greater part of group assignment than negative emotionality. Of interest, both the data-driven behavior (in the five cluster solution) and the brain reserve models (clusters 4 and 5) identify an additional group at low negative emotionality that can be distinguished from other groups on cognitive score.

### 3.2 Reliability of Aging Profiles

We quantitatively assessed the stability of data-driven behavior and brain reserve clustering solutions and their performance relative to a null dataset. ARI, distinctness of clusters (evaluated with silhouette score) and uncertainty of subject probabilities (average entropy) were compared. Higher ARI and silhouette scores indicate better performance, whereas lower entropy indicates better performance.

The brain reserve models achieved high stability relative to null (ARI=0.97; p=0.001), whereas the data-driven behavior model did not outperform the null distribution on stability (ARI=0.98; p=0.169; Fig. 2A & B). Silhouette scores were relatively higher in data-driven behavior models (mean silhouette=0.36, p=0.001; Fig. 2C), compared to brain reserve models (mean silhouette=0.19, p=0.001; Fig. 2D). Entropy scores were lower in data-driven behavior models yet not significantly different from null (mean entropy=0.63, p=0.446; Fig. 2E), whereas brain reserve model entropy was higher overall yet significantly better than null (mean entropy=0.87, p=0.011; Fig. 2F). These significance differences are likely driven by the dimensionality of input features to the clustering algorithm (4 for brain reserve versus 2 for behavior). Overall entropy was relatively lower for participants near profile centroids than for participants near profile boundaries and did not differ between profiles (Supplementary Fig. 4). Despite relatively high entropy suggesting a lack of sparsity in cluster probabilities, the ARI (see above) was also high indicating that the highest-probability cluster was consistent across bootstraps.

**Figure 2.**
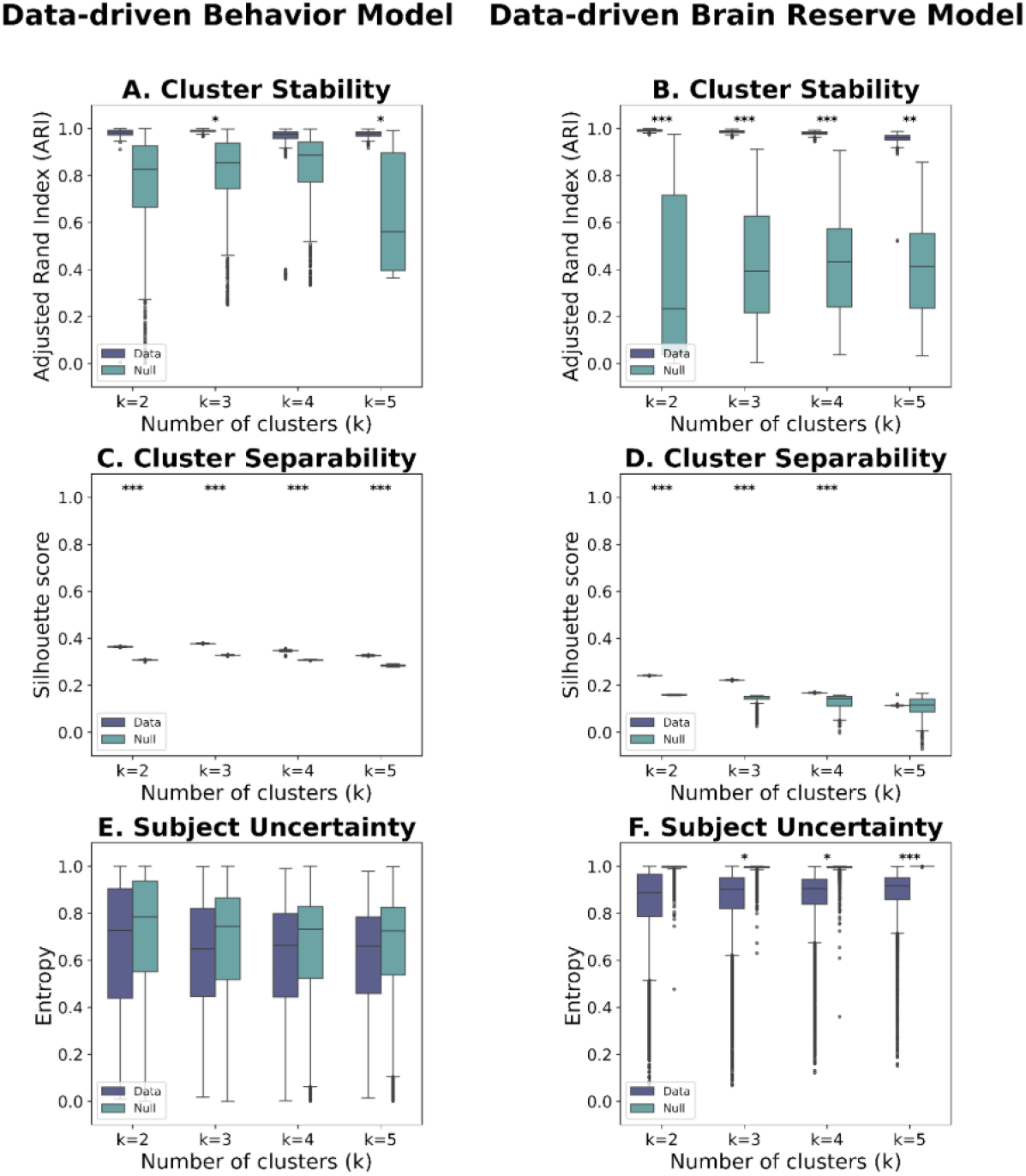
Comparison of cluster solutions for each model against null data. Bootstrapping was used to compare consistency of group assignments for each model. Adjusted Rand Index (higher is better) for the data-driven behavior model (A) and the brain reserve model (B). Silhouette scores (higher is better) for the data-driven behavior model (C) and the brain reserve model (D). Entropy (lower is better) for the data-driven behavior model (E) and the brain reserve model (F). Results reflect 1,000 bootstraps performed using real data (blue) or null data of the same size (green). Significance is indicated with ^*^<0.05, ^**^<0.01, ^***^<0.001.

Overall, results revealed consistency across cluster solutions, without a clear ‘best’ dimensionality. We thus focused on the 4-profile solution in line with the hypotheses outlined in the introduction for the remainder of the manuscript. Complete results across profiles are included in Supplementary Table 4.

### 3.3 Do aging profiles contribute to the emotion paradox?

Cross-sectional relationships between cognition, affect, and age within each profile were considered, to assess how profiles contribute to the emotion paradox. The clinically predefined model had a consistent negative correlation between cognition-age and neuroticism-age across all profiles, consistent with the emotion paradox (i.e., worse cognition with age, yet lower neuroticism with age; Fig. 3A). Similarly, the data-driven behavior model showed a negative relationship between cognition-age and neuroticism-age across profiles (Fig. 3B). The only model that deviated from expected age associations was the brain reserve model (Fig. 3C). Here, neuroticism exhibited a positive correlation with age in profile 1, contrary to the emotion paradox. This profile contained participants with relatively low cognition and low neuroticism (Fig. 2). In brain reserve profile 4,cognition showed no relationship with age, also deviating from the emotion paradox. Profile 4 was composed of subjects with relatively low neuroticism and moderate-high cognition.

**Figure 3.**
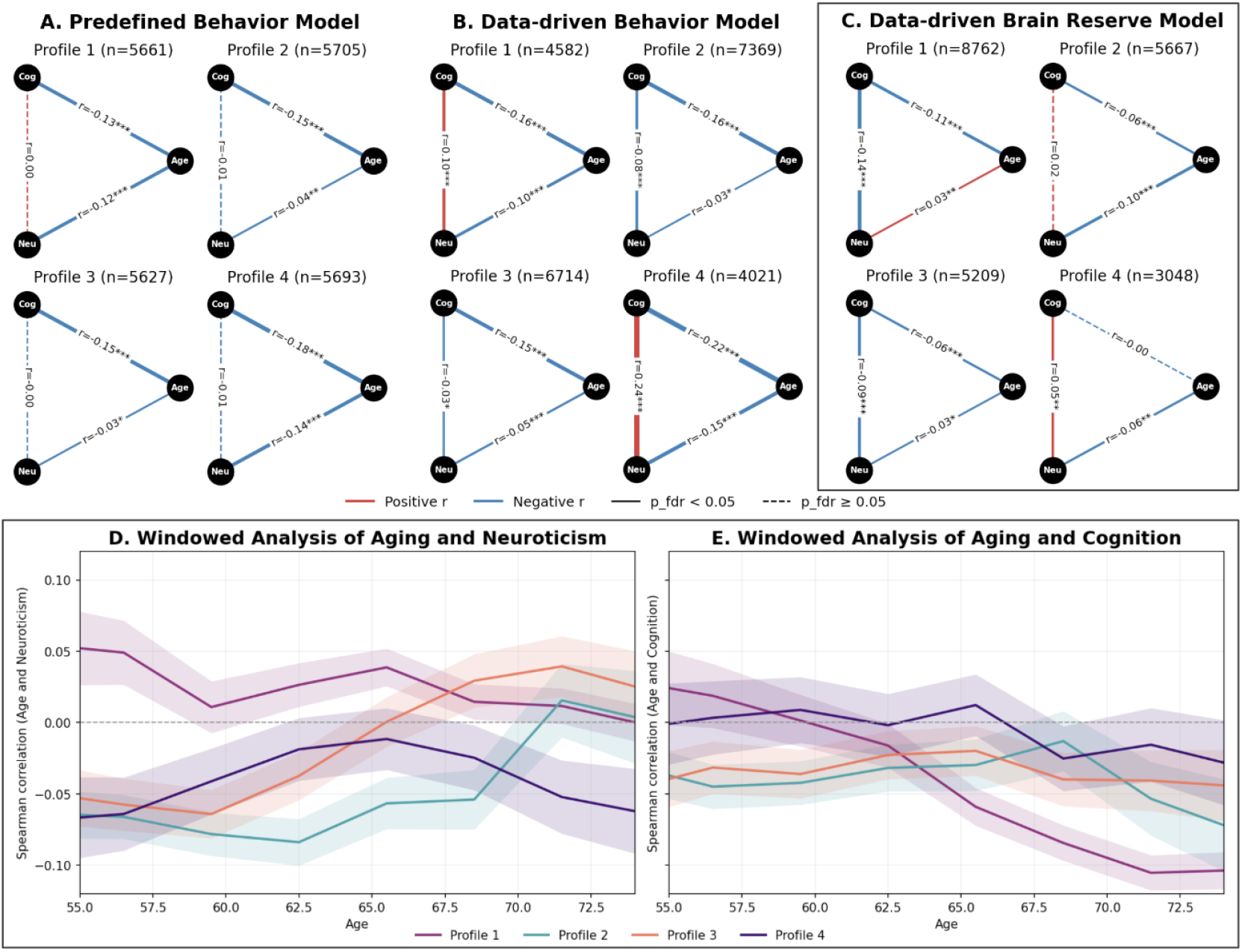
Correlations between age, cognition, and neuroticism for each profile derived from each 4-profile model. A) Correlation between age, cognition, and neuroticism by profile in the predefined model. B) Correlation between age, cognition, and neuroticism by profile in the data-driven behavior model. C) Correlation between age, cognition, and neuroticism by profile in the brain-reserve model (highlighted by border because the remainder of the manuscript focuses on these profiles). Lower hand plots show follow-up analyses for the brain reserve model (C) to visualize age-neuroticism (D) and age-cognition (E) associations broken down into cross-sectional age windows of 15 years, sliding by 3 years. Results for each year are shown for the window centroid, line colors refer to profiles 1-4 derived from the brain reserve model, and shaded regions depict the standard error of mean. In plots A-C, blue lines indicate negative correlations, red lines indicate positive correlations, and line thickness reflects correlation strength. Significance is indicated with ^*^<0.05,^**^<0.01, ^***^<0.001, and non-significant correlations are indicated by dashed lines. Cog = cognition, Neu = neuroticism.

We leveraged a windowed analysis to assess whether associations between age and cognition/neuroticism were stable across aging for brain reserve model profiles. The results (Fig 3D, 3E, Supplementary Fig. 5) revealed consistent positive associations between age and neuroticism in profile 1. On the other hand, the age-neuroticism association showed a tipping point in profile 3 around ∼60 years old and in profile 2 the tipping point came later (∼67 years old), consistent with profile 2 being healthier overall than profile 3 (Supplementary Fig. 5A). Associations between age and cognition were relatively stable across age for profiles 2-4, but suggested increasingly rapid cognitive decline with age for profile 1, suggesting poor health outlook in this profile (Fig. 3E).

Taken together, the brain reserve model was the only model to suggest the existence of differential aging profiles that both confirmed the previously reported emotion paradox (across 48% of participants in profiles 2 and 3), while also refuting the emotion paradox in a substantial subset of individuals (52% of participants in profiles 1 and 4). Based on these findings, we focused on neuroanatomical differences and validation results in the brain-reserve model for subsequent sections.

### 3.4 Volumetric differences between aging profiles

To assess neuroanatomical characteristics of the aging profiles, neural volumes were compared between brain reserve profiles. Profile 1, the group exhibiting the lowest cognition and low-to-moderate negative emotionality (Fig. 1), had broadly lower brain volumes than the population average (Fig. 4), whereas profile 2, the group exhibiting the highest cognition with low-moderate negative emotionality (Fig. 1), had on average larger brain volumes than the population average (Fig. 4). Profiles 3 and 4 were closer to average in most regions (Fig. 4).

**Figure 4.**
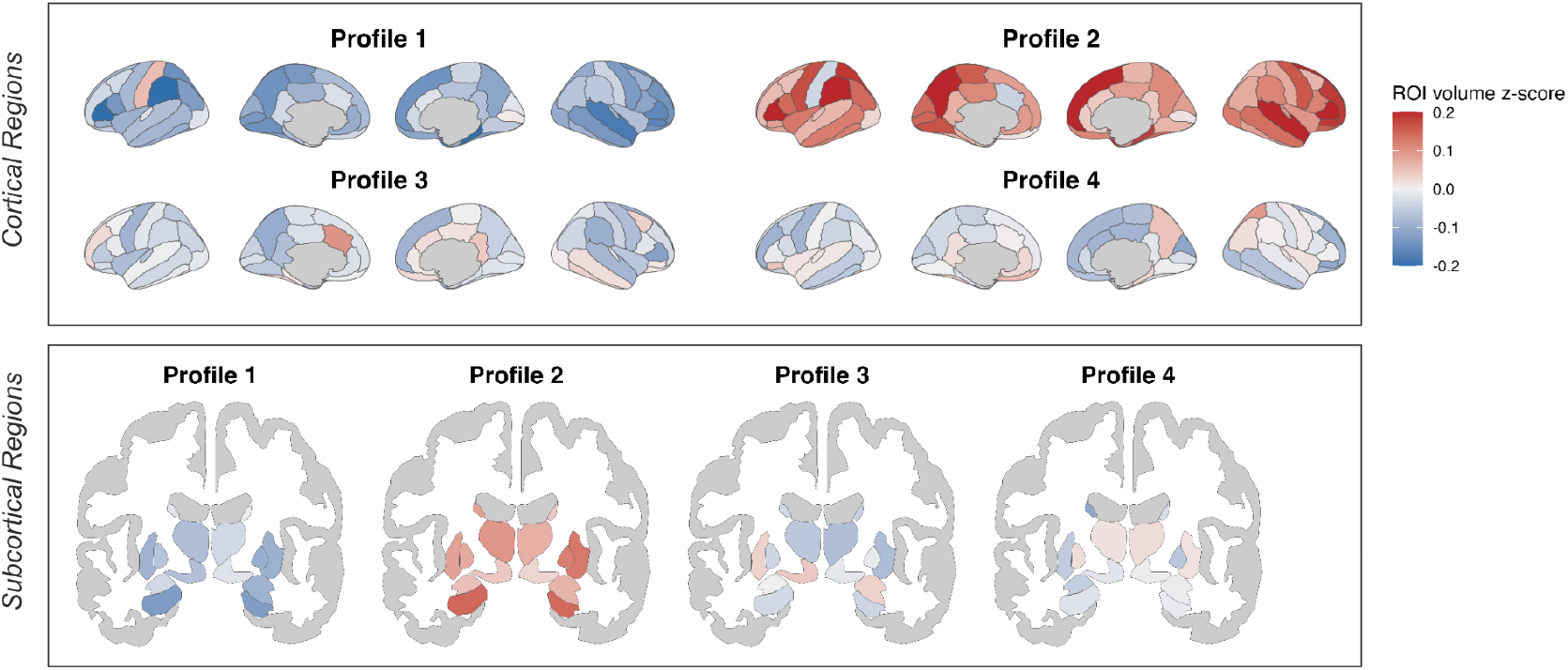
Volume differences by profile in the brain reserve model relative to population average. Gray matter volumes in each brain region were z-scored across the full dataset and then averaged within each profile to reveal neuroanatomical characteristics. Notably, the brain reserve model incorporated brain volumes as part of the CCA, so the presence of volume differences between aging profiles is expected. As such, these results are intended purely to provide descriptive context.

### 3.5 Validation of aging profiles

To validate the aging profiles, we assessed differences on demographic and etiological measures that were not used for profile definition. Statistical comparisons were performed for all profile solutions (supplementary Table 5). The data-driven brain reserve model (mean Ω=0.07) outperformed other models (mean Ω=0.02) in the validation results and is the focus of the description below.

There was a large main effect of age (Ω=0.23, p<1^*^10^-300^), which was consistent across profile dimensionalities. Profile 1 was significantly older than other profiles (mean age=68.30; Fig. 5A), consistent with relatively lower cognition scores and reduced gray matter volume. Conversely, profile 2 was significantly younger than other profiles (mean age=59.42; Fig. 5A), consistent with relatively higher cognition scores and larger gray matter volume.

There was a moderate main effect of sex (Ω=0.12, p<1^*^10^-300^), which was consistent across profile dimensionalities. Profiles 1 and 3 were predominantly female (61.65% and 69.40% female respectively), whereas profile 2 was predominantly male (74.57% male) and profile 4 was the most balanced (56.86% male) (Fig. 5B). These sex differences may partly explain the brain size differences between profiles both at the regional level (Fig. 4) and the large main effect of total intracranial volume (main effect Ω=0.25, p<1^*^10^-300^; Fig. 5C).

**Figure 5.**
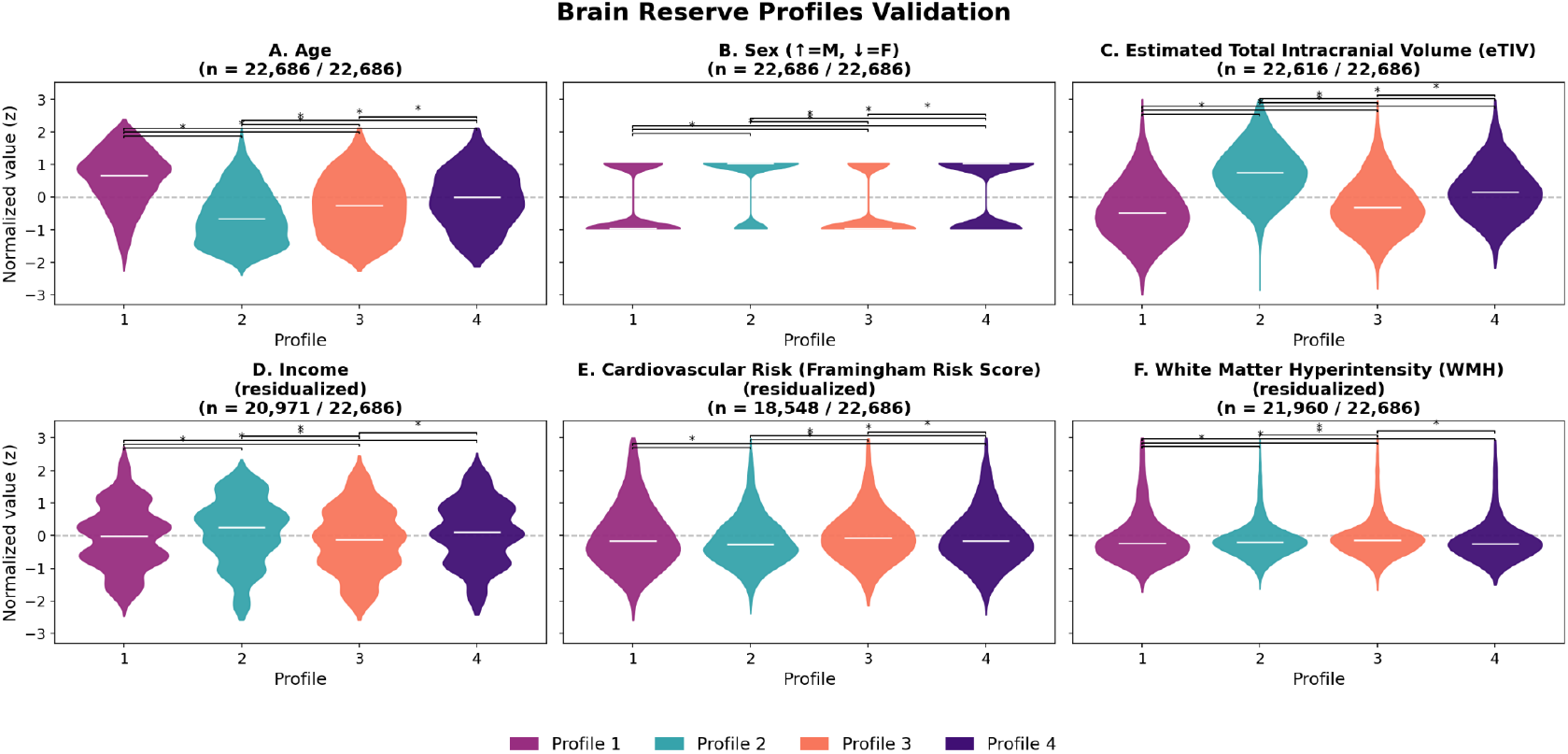
Differences Between Aging Profiles. Significant ANOVA and ANCOVA results comparing profiles derived from the brain reserve model. Y axes show z-scored values and x axes list the profiles (1-4). Full statistical results for all models can be found in supplementary table 5. Differences between profiles visualized by black lines were identified using post-hoc Tukey tests. A) Comparison of age (based on ANOVA). B) Comparison of sex based on ANOVA (-1 = female and 1 = male). C) Comparison of total intracranial volume (based on ANOVA). D) Comparison of income (based on ANCOVA controlled for age and sex). The Y axis shows values that were z-scored and then residualized. Income was reported as a categorical variable with higher categories indicating a higher income bracket. E) Comparison of Framingham risk score (based on ANCOVA controlled for age and sex). F) Comparison of white matter hyperintensity volume (based on ANCOVA controlled for age, sex, and total intracranial volume). The Y axis shows values that were z-scored and then residualized. Sample sizes for each variable are shown at the top out of a maximum sample of N=22,686 and vary based on data availability.

For other demographic variables, there was a small main effect of income (Ω=0.04, p=8.3^*^10^-222^), which persisted after correcting for age and sex (Ω=0.013, p=2.84^*^10^-51^). Here, profile 2 included proportionally more high-earners than the other profiles (Fig. 5D). The effect size of education was minimal yet significant due to high power (Ω=0.003, p=2.1^*^10^-15^).

In terms of cardiovascular risk, there was a moderate main effect of Framingham scores (Ω=0.07, p=2.6^*^10^-291^), with relatively higher Framingham risk scores in profile 3 (Fig. 5E). Similarly, there was a small main effect for total white matter hyperintensity volume (Ω=0.04, p=2.3^*^10^-202^), which survived correction for age, sex, and total intracranial volume (Ω=0.01, p=5.0^*^10^-51^; Fig. 5F). Consistent with the Framingham results, profile 3 had relatively higher white matter hyperintensity volume. The effect size of metabolic equivalent activity scores was minimal yet significant due to high power (Ω=0.004, p=9.2^*^10^-19^).

In terms of inflammatory markers, there was a small main effect of GFAP (Ω=0.02, p=0.0003), which did not survive correction for age and sex. The effect size of c-reactive protein was minimal yet significant due to high power (Ω=0.002, p=1.5^*^10^-11^).

Polygenic risk score for Alzheimer’s disease did not differ between profiles (Ω=0, p=0.99). Although, our sample size was relatively small (N=4,033), since at the time of publication this data was available for a minority of subjects.

## Discussion

Our goal was to identify possible profiles of aging using cognitive, affective, and brain reserve data in a large population of older adults. We hypothesized that aging profiles may offer an explanation for contradictory relationships between age, cognition, and mental health. Despite the inverse association between emotional wellbeing [2] and declines in cognition [1] (described as the emotion paradox [4]), poor mental health in older age is linked to a three-fold increased risk of dementia [5–7]. Our findings supported the existence of differential aging profiles. We focused on four profiles derived from a brain reserve model because it consistently achieved the strongest profile-differentiation in validation results and because it helps explain contradictory relationships between age and cognitive/mental health. These findings highlight the importance of a nuanced approach to aging. In the remainder of this discussion, we summarize the four brain reserve profiles of aging to contextualize our findings.

Profile 1 mapped onto our hypothesized suboptimal aging profile as characterized by relatively lower cognitive scores and relatively higher neuroticism scores. This was the oldest profile, included the highest proportion of women, and had the lowest brain volume. Notably, profile 1 diverged from the emotion paradox in that older age was associated with worsening (rather than improved) emotional wellbeing. These results expand on prior work that showed increased frequency of positive emotions minus negative emotions from 20-50 years old, which leveled off between 50 and 70, and declined after 70 [32]. Our findings revealed consistent increases in negative emotionality at age 50 in a subset of older aged individuals, and increasingly steep declines in cognition with age. Notably, profile 1 had the highest proportion of low earners (yet no difference in education), suggesting this profile was largely made up of non-working or post-retirement participants. Prior work has shown high heterogeneity in the relationship between retirement and affect [33], with evidence of increased post-retirement depression [34], especially in those with positive work identification [33]. These results suggest this suboptimal aging profile could reflect a pre-dementia trajectory suffering from many risk factors that are potentially modifiable [35]. Without intervention, the chronic stress in profile 1 may accelerate neuronal loss leading to pathological aging such as neurodegeneration [36]. Therefore, profile 1 may benefit from preventive strategies to mitigate progression [37], which could impact population health outcomes given that profile 1 was the largest of all profiles, containing 39% of all participants.

Profile 2 was associated with higher cognitive ability and lower neuroticism than the dataset average. Profile 2 had the largest fraction of males, which likely contributed to this group’s relatively larger intracranial volume. As the youngest profile, profile 2 likely had the greatest fraction of individuals below retirement age. Notably, profile 2 had the most participants in high income brackets. Prior work has shown associations between higher socioeconomic status and more positive aging outcomes in both cognitive and mental health [38] and more positive aging perception [39], consistent with our findings in profile 2. Nevertheless, profile 2 had relationships between age-cognition and age-neuroticism consistent with the emotion paradox. The presence of the aging paradox in this younger profile is consistent with prior work showing improvements in emotional wellbeing with age up until 50 [32]. Given that profile 2 was younger than other profiles, it is possible that participants may move into other aging profiles with age. Future work leveraging rich longitudinal data will be needed to assess this possibility.

Unlike the others, profile 3 was not associated with a particular cognitive range. Instead it was associated with high neuroticism scores irrespective of cognition. Profile 3 was younger than all profiles but profile 2, and like profile 1 was predominantly female. Younger age and female gender have been associated with lower risk for and incidence of heart disease, though sex differences in lifetime cardiovascular risk have been overstated [40–42]. Profile 3 had higher Framingham risk scores and white matter hyperintensity than other profiles after controlling for age and sex. White matter hyperintensity is an important link between cardiovascular risk and dementia [43], detectable before overt cognitive impairment manifests [44]. Women are more likely to have white matter hyperintensities than men [44–46], consistent with the demographics of profile 3. Framingham risk score is a clinical tool used to estimate likelihood of future cardiovascular events [47]. Given the elevated Framingham risk score and volume of white matter hyperintensities, profile 3 may represent high cardiovascular risk individuals. Cardiovascular risk has shown a dose–response association with depression [48], and risk of vascular dementia was also shown to increase monotonically with greater negative affective symptoms [49,50]. As profile 3 represented both high affective symptoms and high cardiovascular risk, we would predict that individuals in profile 3 may have greater risk of vascular dementia or other cardiovascular events than other profiles in older age.

Profile 4 evidenced average cognitive ability and low neuroticism. Prior work has indicated psychological resilience moderates epigenetic aging [51]. Similarly, low neuroticism has been linked to reduced frailty [52] and to preserved telomere length [53] in aging. As such, results in profile 4 were consistent with work suggesting low levels of neuroticism may slow aging. Profile 4’s demographics were close to average on age, income, brain volume, and were balanced between male and female subjects. Similarly, profile 4 showed cardiovascular factors close to the sample average. Interestingly, profile 4 was the only group that did not show a cross-sectional decline in cognition with age, which persisted into older age windows. This finding of stable cognition suggests members of profile 4 may be less vulnerable to cognitive decline than our other participants, analogous to resilient/slow decline groups identified in other studies [54,55]. As the smallest profile, profile 4 may represent the 13% of older-aged people with a resilient aging trajectory, and we hypothesize that these participants may develop dementia at a lower rate than members of other profiles.

Although this paper revealed new insights into potentially diverging aging profiles, there are limitations to consider. Most importantly, this is a cross sectional study. The conclusions we can draw about aging within individuals are limited. Future work is needed to test longitudinal outcomes of aging profiles. We are furthermore limited in measures used to assess cognition in our sample. Although summary scores have been shown to provide insight into cognitive abilities [24], a comprehensive cognitive battery would provide richer information about participant’s cognition. Furthermore, the UK Biobank represents a relatively higher socioeconomic status cohort that is majority White [56]. Given our findings linked to income in profile 2, further work in diverse populations is needed. Lastly, although our results provide evidence for - and insights from - stratified aging profiles, our findings did not resolve the optimal number of profiles. An explanation for the consistency of findings across profile dimensions is the presence of a hierarchy with multiple potential levels of profile differentiation (e.g., a single suboptimal aging group may further split into cardiovascular risk versus Alzheimer risk, etc.). Even though this hierarchy can be meaningfully cut off at multiple levels, future work may wish to explore dimensionalities of aging profiles that best inform clinical decision making and/or preventative strategies.

In conclusion, our results revealed differentiable profiles of aging that may explain paradoxical links between cognition and emotional wellbeing in older age. These findings highlight the importance of a nuanced approach to understanding aging, and highlight opportunities for targeted intervention to improve healthspan in older age [57,58].

## Data sharing statement

UKB data are available following an access application process: https://www.ukbiobank.ac.uk/enable-your-research/apply-for-access. This research was performed under UK Biobank application number 47267.

## Code sharing statement

Code for this proposal are available on GitHub: https://github.com/PersonomicsLab/AgingProfile_BrainReserve

## Author contributions

Conceptualization: SJ, LS, KT, JB

Data Curation: SJ, LS, PL

Formal Analysis: SJ, LS, KT, BF, SR, XL, HM, KH, FA, TE

Funding Acquisition: JB

Methodology: SJ, LS, SR, TE, JB

Supervision: JB

Visualization: SJ

Writing – Original Draft: LS

Writing – Review & Editing: SJ, LS, BF, KH, FA, JB

