## Supplemental Material for "Profiles of Aging Based on Cognition, Affect, and Brain Reserve"

### Supplementary Materials

**Supplementary Table 1**. Brain reserve variables

| **Gray matter volume UK Biobank variable IDs** |
|---|
| Accumbens-area (26564 & 26595) |
| Amygdala (26563 & 26594) |
| Brainstem (26526) |
| Caudal anterior cingulate (27205 & 27298) |
| Caudal middle frontal (27206 & 27299) |
| Caudate (26559 & 26590) |
| Cerebellum-Cortex (26557 & 26588) |
| Cuneus (27207 & 27300) |
| Entorhinal (27208 & 27301) |
| Fusiform (27209 & 27302) |
| Hippocampus (26562 & 26593) |
| Inferior parietal (27210 & 27303) |
| Inferior temporal (27211 & 27304) |
| Insula (27235 & 27328) |
| Isthmus cingulate (27212 & 27305) |
| Lateral occipital (27213 & 27306) |
| Lateral orbitofrontal (27214 & 27307) |
| Lingual (27215 & 27308) |
| Medial orbitofrontal (27216 & 27309) |
| Middle temporal (27217 & 27310) |
| Pallidum (26561 & 26592) |
| Paracentral (27219 & 27312) |
| Parahippocampal (27218 & 27311) |
| Pars opercularis (27220 & 27313) |
| Pars orbitalis (27221 & 27314) |
| Pars triangularis (27222 & 27315) |
| Pericalcarine (27223 & 27316) |
| Postcentral (27224 & 27317) |
| Posteriorcingulate (27225 & 27318) |
| Precentral (27226 & 27319) |
| Precuneus (27227 & 27320) |
| Putamen (26560 & 26591) |
| Rostralanteriorcingulate (27228 & 27321) |
| Rostralmiddlefrontal (27229 & 27322) |
| Superiorfrontal (27230 & 27323) |
| Superiorparietal (27231 & 27324) |
| Superiortemporal(27232 & 27325) |
| Supramarginal (27233 & 27326) |
| Thalamus (26558 & 26589) |
| Transversetemporal (27234 & 27327) |
| VentralDC (26565 & 26596) |

**Supplementary Table 2**. Validation measures. The Framingham Risk Score was calculated from the UK Biobank as described in previous work [1]. For the Framingham Risk Score calculation, antihypertensive use was identified using medication codes found in Supplementary Table 4 of [1].

| Domain | Variable | UKB Variable ID |
|---|---|---|
| Demographics | Age<br>Sex<br>Income<br>Education | 21003<br>31<br>738<br>26414 |
| Brain size | Total intracranial volume | 26521 |
| Cardiovascular health | Framingham Risk Score calculated as described in [1] based on the following variables | Age: 21003<br>Antihypertensive use: 20003<br>Diabetes status: 41270, 41202, 41204<br>HDL: 30760<br>Sex: 31<br>Smoking status: 20116<br>Systolic blood pressure: 93, 4080<br>Total cholesterol: 30690 |
| | Summed metabolic equivalents (MET) minutes per week for all activity | 22040 |
| | White matter hyper intensities | 25781 |
| Inflammation | C-reactive protein | 30712 |
| | Glial fibrillary acidic protein (GFAP) | 31042 |
| Risk for Alzheimer's Disease | Polygenic risk score for Alzheimer's Disease | 26207 |

**Supplementary Figure 1.** Group assignment by cognition and neuroticism for different numbers of clusters (k). A) Data-driven behavior group assignments at k=2. Group assignment is primarily influenced by negative emotionality. B) Brain reserve group assignment where k=2. Group assignment is primarily influenced by cognitive score. C) Data-driven behavior group assignment at k=3. D) Brain reserve group assignment where k = 3. E) Data-driven behavior group assignment based on clinical measures where k = 4. F) CCA group assignment where k = 4. G) Group assignment based on clinical measures where k=5. H) CCA group assignment where k=5. I) Group assignments based on predefined thresholds.

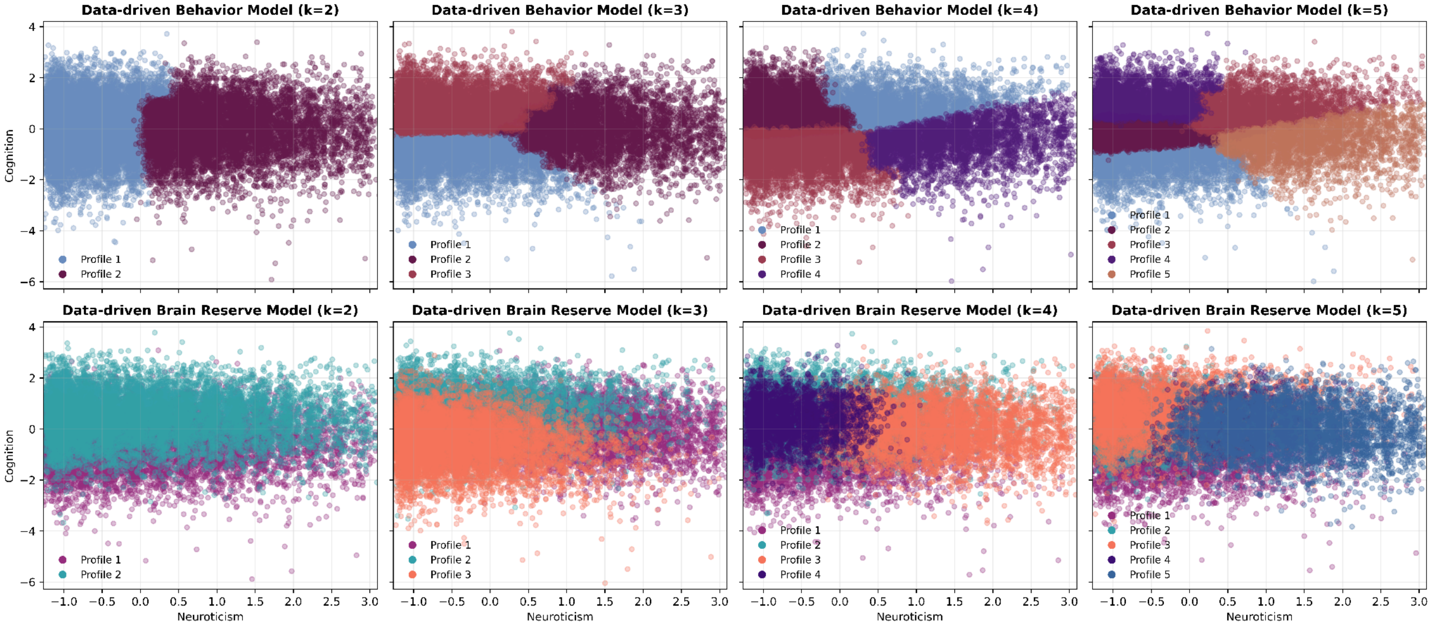

**Supplementary Figure 2.** Adjusted Rand Index between 4 cluster solutions.

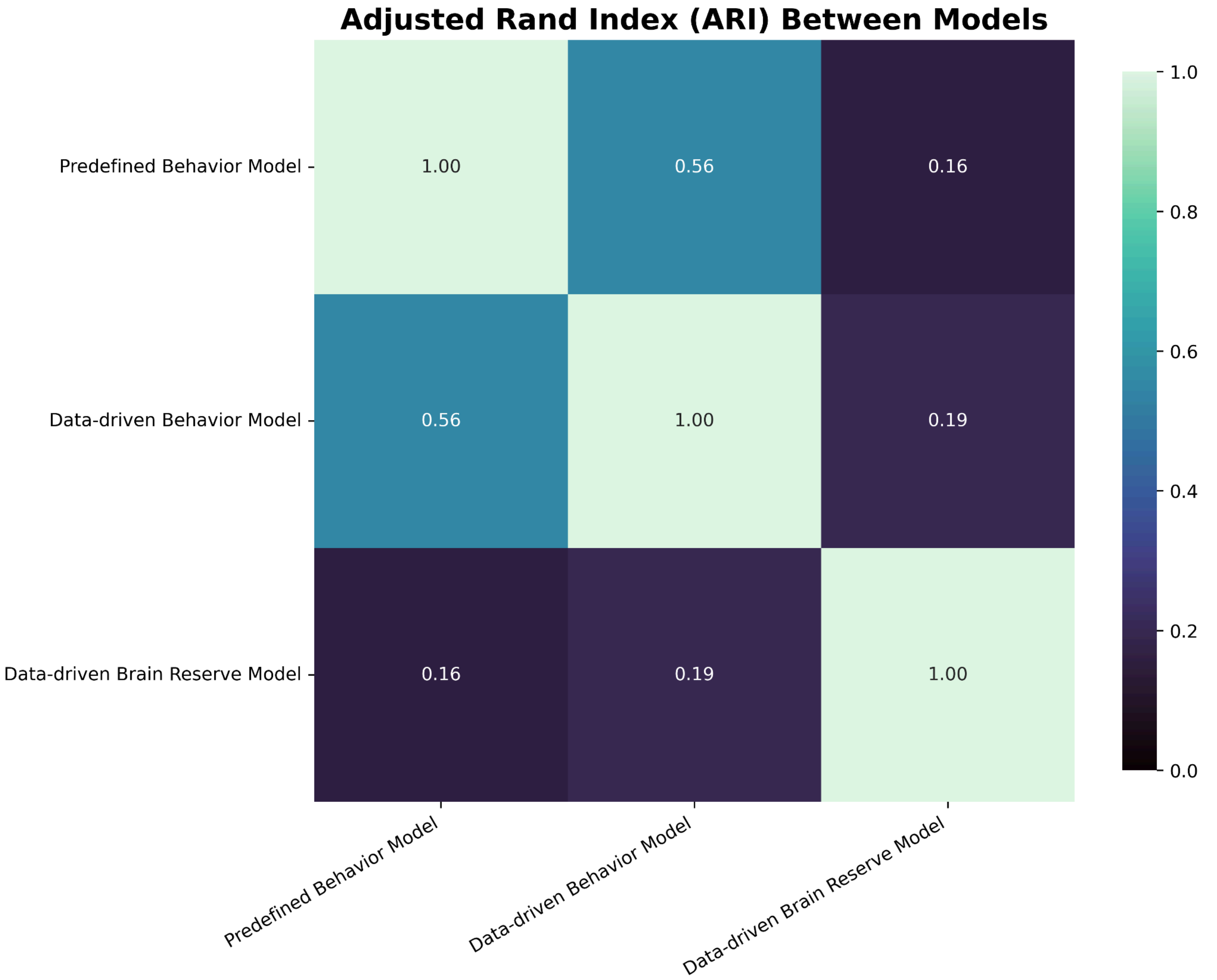

**Supplementary Figure 3.** Brain reserve 4-profile solution shown in the space of behavioral canonical covariates (left) and in the space of brain canonical covariates (right), as a function of cognition, neuroticism, age, and age.

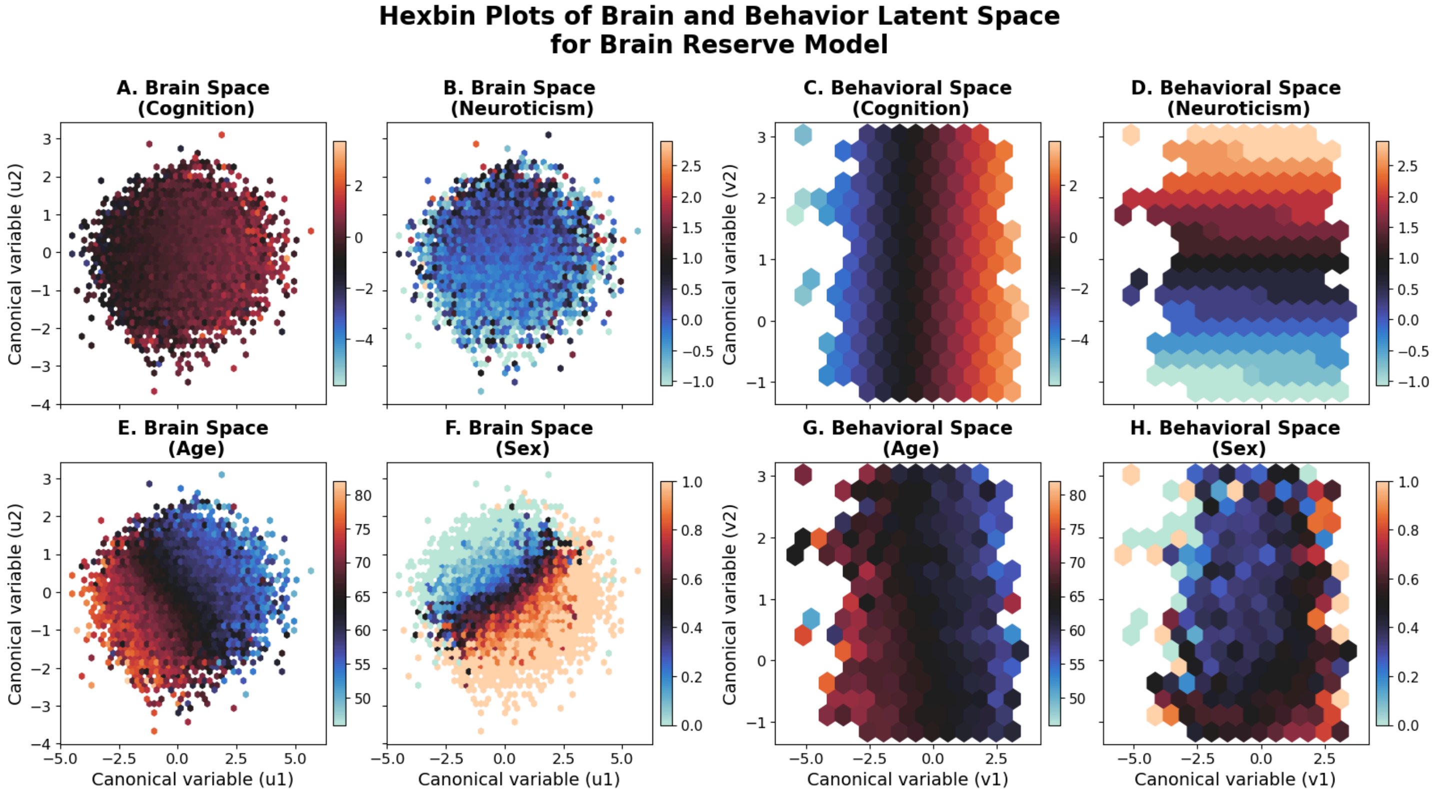

**Supplementary Table 3**. Descriptive statistics for the four aging profiles derived using the brain reserve model.

| Profile | Size (n) | Age (mean) | Age (std) | Percentage of Female | Percentage of Male | eTIV (mean) | eTIV (std) | Income (median) | Income (IQR) | Framingham Risk Score (mean) | Framingham Risk Score (std) | WMH (mean) | WMH (std) |
|---|---|---|---|---|---|---|---|---|---|---|---|---|---|
| 1 | 8762 | 68.3 | 6.53 | 61.65 | 38.35 | 1490991 | 139145 | 3 | 1 | 28.77 | 15.64 | 6851.69 | 8316.57 |
| 2 | 5667 | 59.42 | 6.37 | 25.43 | 74.57 | 1675064 | 131394 | 3 | 2 | 20.47 | 11.75 | 3743.53 | 4656.27 |
| 3 | 5209 | 62.1 | 6.94 | 69.4 | 30.6 | 1515983 | 127912 | 3 | 2 | 20.51 | 13.4 | 4297.53 | 5279.85 |
| 4 | 3048 | 63.43 | 6.69 | 43.14 | 56.86 | 1584485 | 126516 | 3 | 2 | 24.27 | 14.23 | 4567.68 | 5477.39 |

**Supplementary Figure 4.** Entropy results for the Brain reserve 4-profile solution, shown in brain canonical covariate space (A), behavioral canonical covariate space (B), and separate per profile (C).

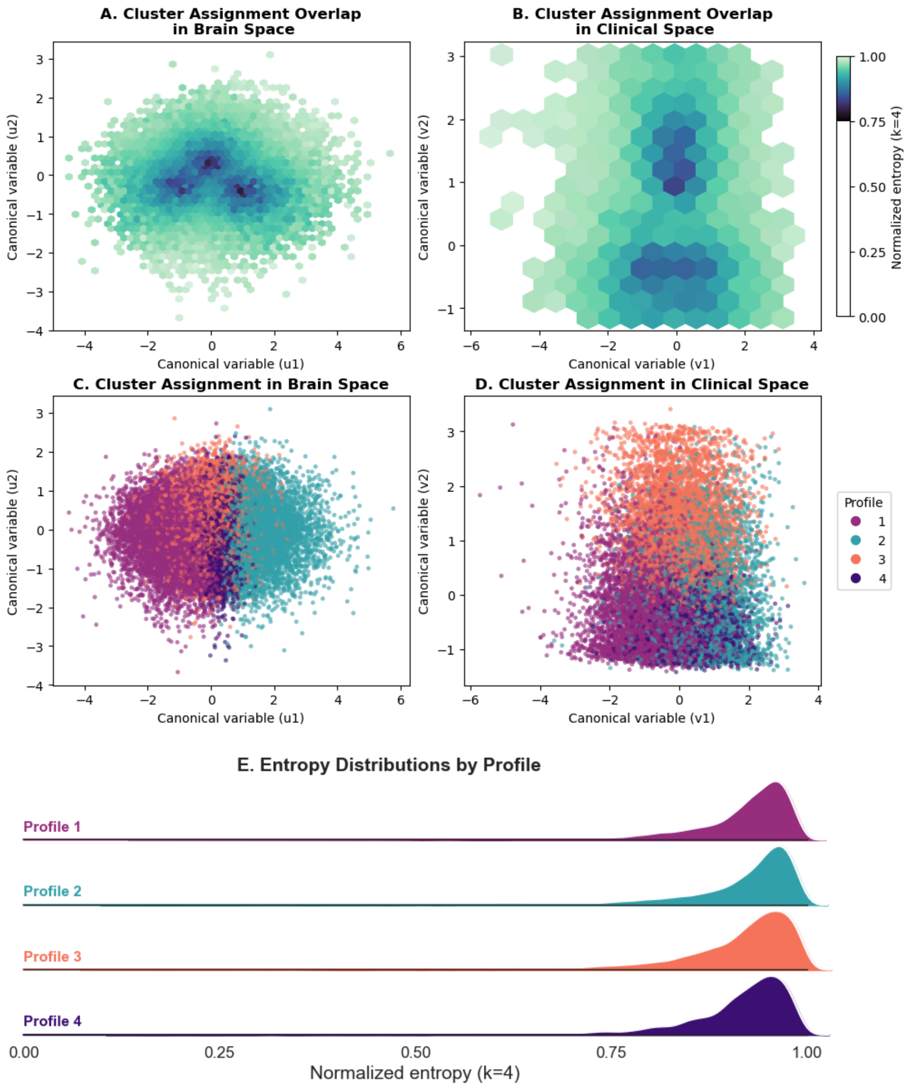

**Supplementary Table 4**. Reliability results for all data-driven profiles from both the behavior and the brain reserve models.

| Metric | Model | k=2 | k=3 | k=4 | k=5 |
|---|---|---|---|---|---|
| **ARI** | Data-driven behavior | 0.075 | 0.032 | 0.169 | 0.028 |
| | Data-driven brain reserve | 0.001 | 0.001 | 0.001 | 0.001 |
| **Silhouette score** | Data-driven behavior | 0.001 | 0.001 | 0.001 | 0.001 |
| | Data-driven brain reserve | 0.001 | 0.001 | 0.001 | 0.525 |
| **Entropy** | Data-driven behavior | 0.439 | 0.431 | 0.446 | 0.426 |
| | Data-driven brain reserve | 0.068 | 0.027 | 0.011 | 0.001 |

**Supplementary Figure 5.** Windowed associations for the 4-profile brain reserve model. Associations between age and neuroticism (A) and age and cognition (B) displayed separate for each of the 4 profiles. Each point reflects the correlation estimated across a 15-year window, where the x-axis reflects the window mean. Sample size is displayed on the y axis.

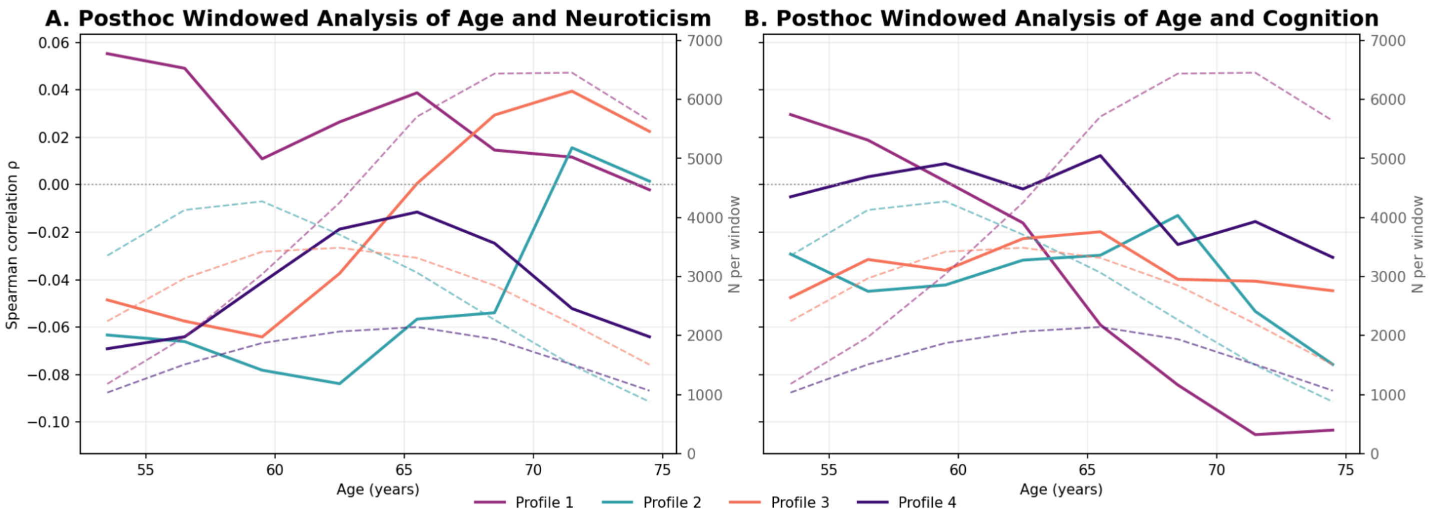

**Supplementary Table 5**. Results of all ANOVA and ANCOVA validation analyses are provided as a separate file entitled ‘Supplementary_Table_5.xlsx
